# Type 2 Diabetes Risk Alleles Reveal a Role for Peptidylglycine Alpha-amidating Monooxygenase in Beta Cell Function

**DOI:** 10.1101/158642

**Authors:** Anne Raimondo, Soren K. Thomsen, Benoit Hastoy, Mahesh M. Umapathysivam, Xiao-Qing Dai, Jocelyn E Manning Fox, Amy Barrett, Christopher J. Groves, Austin Bautista, Nicola L. Beer, Anne Clark, Patrick E. MacDonald, Patrik Rorsman, Anna L. Gloyn

**Author notes:** Contact Information: Anna L. Gloyn, Oxford Centre for Diabetes, Endocrinology & Metabolism, University of Oxford, Churchill Hospital, Headington OX3 7LE. These authors contributed equally to this work (Anne Raimondo, Soren K. Thomsen).

## Abstract

Molecular mechanisms underpinning the genetic risk for type 2 diabetes (T2D) remain poorly understood, hindering translation into new therapies. Recently, genome-wide studies identified two coding variants in *Peptidylglycine Alpha-amidating Monooxygenase* (*PAM*) associated with T2D risk and measures of beta cell dysfunction. Here, we demonstrate that both risk alleles impact negatively on overall PAM activity, but via distinct effects on expression and catalytic function. In a human beta cell model, *PAM* silencing caused decreased insulin content and altered dynamics of granule exocytosis. Analysis of primary human beta cells from cadaveric donors confirmed an effect on exocytosis in carriers of the p.D563G T2D-risk allele. Finally, we show that the granular packaging protein Chromogranin A is a PAM substrate and a strong candidate for mediating downstream effects on insulin secretion. Taken together, our results establish a role for PAM in beta cell function, and uncover a novel mechanism for T2D-associated *PAM* alleles.

## INTRODUCTION

Human genetics has the potential to identify novel mechanisms for disease, which in turn present opportunities for clinical translation. Over the past 10 years, genome-wide association studies (GWAS) have identified over 150 genomic regions robustly associated with type 2 diabetes (T2D) risk, and have revealed a central role for pancreatic beta cell dysfunction in T2D pathogenesis (1–3). However, the translation of GWAS signals into molecular mechanisms for disease has been slow, primarily due to uncertainty over the transcripts through which non-coding association signals operate (4). Recently, studies focused on identifying coding variant association signals have identified nonsynonymous alleles in *Peptidylglycine Alpha-amidating Monooxygenase (PAM)* independently associated with T2D risk (rs78408340, p.S539W; and rs35658696, p.D563G), providing a direct link to the effector transcript (3, 5). The T2D risk alleles are also associated with reduced insulinogenic index – a measure of glucose-stimulated insulin secretion – suggesting that their effects are mediated via altered beta cell function (5–7).

*PAM* encodes peptidylglycine alpha-amidating monooxygenase (PAM), an enzyme in neuroendocrine cells that modifies peptides with a C-terminal glycine to create peptide-amides (8–10). Amidation can dramatically increase the biological potency of a peptide relative to its unmodified glycine-extended conjugate (11). PAM is localised to the Golgi, where it is packaged with other endocrine proteins into nascent granules (8, 12). The functional enzyme exists in both integral membrane and luminal forms, the latter of which is co-secreted with the endocrine peptide(s) (8, 10, 13).

Despite reports of PAM expression in pancreatic islets, a functional role in beta cells has not yet been described (14). Insulin itself is not a PAM substrate, so the effects of *PAM* on beta cell function are therefore mediated by other peptide(s). We hypothesised that T2D-associated *PAM* missense alleles reduce PAM function, affecting amidation of peptides critical for insulin secretion. We demonstrate that both diabetes risk alleles negatively impact on PAM expression and/or activity, and elucidate an endogenous role for PAM in insulin granule packaging and release from beta cells. We also show that PAM amidates the granule packaging factor Chromogranin A (CgA), and establish this neuroendocrine peptide as a likely downstream effector of PAM in beta cells. Our results are consistent with the direction and magnitude of effects for T2D-associated risk alleles in *PAM*, and establish molecular mechanisms for their impact on disease susceptibility.

## RESULTS

### T2D-associated PAM alleles cause *PAM* loss-of-function

PAM is a bifunctional enzyme, possessing two contiguous catalytic domains: peptidylglycine alpha-hydroxylating monooxygenase (PHM), and peptidyl alpha-hydroxyglycine alpha-amidating lyase (PAL) (10). The p.S539W and p.D563G missense mutations are both located within the PAL domain and are predicted by *in silico* tools (SIFT, PolyPhen2) to be damaging, suggesting that they could affect enzymatic activity. To test this, we generated Human Embryonic Kidney (HEK) 293 cell lines expressing recombinant secreted (nonintegral membrane) PAM protein for *in vitro* amidation assays. In line with previous observations, PAM was constitutively released into supernatant (Supplementary Figure 1A) (15). WT-PAM and D563G-PAM were robustly produced, as well as an additional catalytically inactive mutant protein, Y651F-PAM, which was used as a control (16). Interestingly, we were unable to detect S539W-PAM expression (Supplementary Figure 1A). This was observed across three independently derived cell lines, and was not due to cellular retention of S539W-PAM (data not shown).

We subsequently developed a cell-free kinetic assay capable of measuring PAM amidating activity via spectrophotometric detection of converted glyoxylate, a by-product of the amidation reaction (17, 18). Matching each reaction for PAM input, we observed reduced amidating activity for D563G-PAM (p=1.0x10^-5^) and Y651F-PAM (p=4.1x10^-6^) (Figure 1A). In agreement with its lack of expression, supernatant from S539W-PAM-transfected cells was inactive in this assay (Supplementary Figure 1B). Further analysis showed no significant difference in substrate affinity between WT-PAM and D563G-PAM (Km 0.95mmol/L vs 1.02mmol/L, p=0.44), suggesting that the p.D563G substitution affects K_cat_ (Supplementary Figure 1C). These results demonstrate that the T2D-associated *PAM* coding alleles decrease PAM function via a combination of defective expression and/or reduced catalysis.

**Figure 1.**
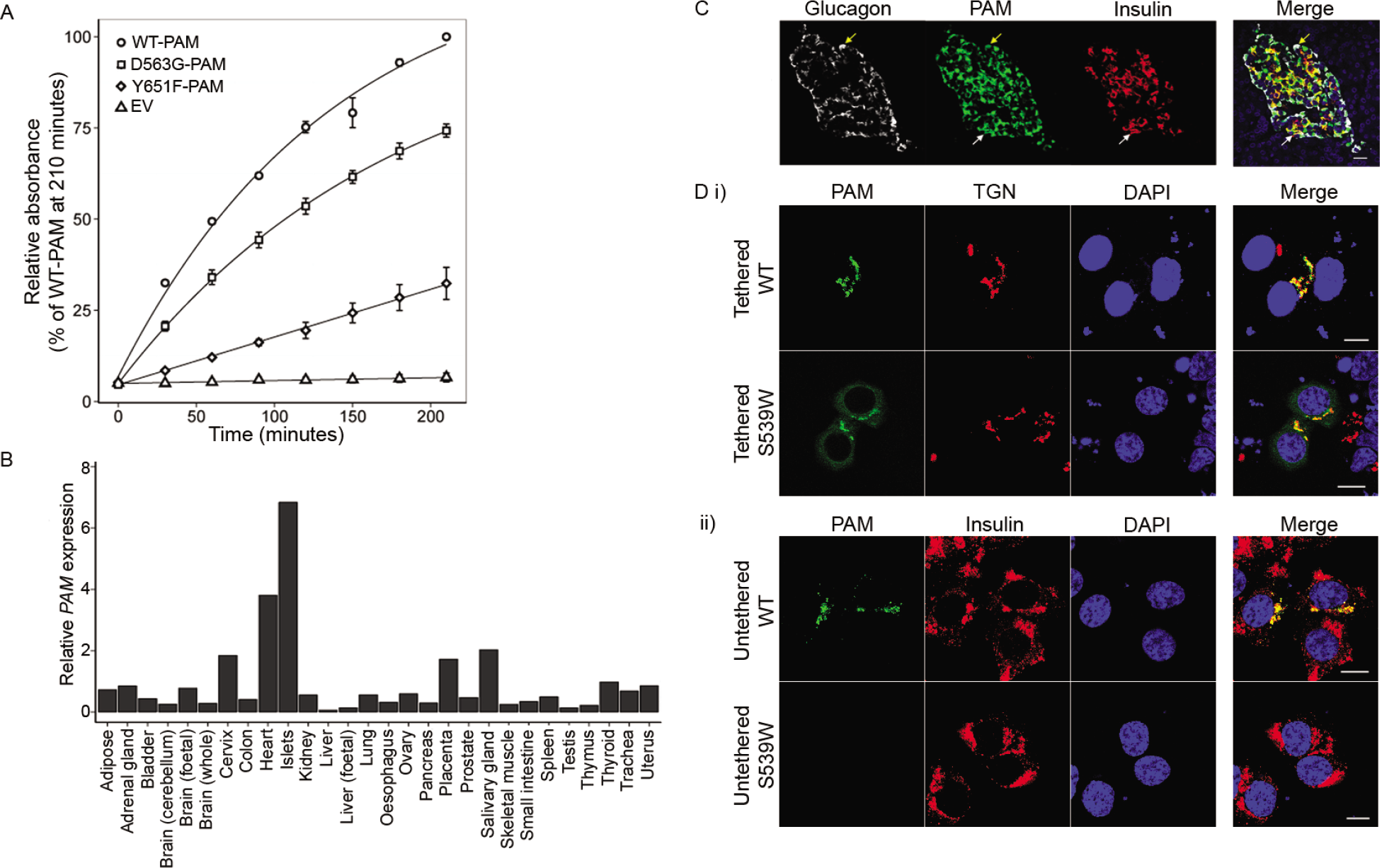
Analysis of WT and variant PAM function and expression. A) Amidating activity of WT-PAM (circles), D563G-PAM (squares), Y651F-PAM (diamonds), or empty vector (EV) (triangles) *in vitro* (n=4). Error bars are mean ±SEM. B) *PAM* expression in human tissues (n=1). C) Endogenous glucagon (white), PAM (green), and insulin (red) expression in human pancreas. White and yellow arrows indicate co-localisation of PAM with insulin and glucagon respectively. Scale bar, 20μm. D) EndoC-0H1 cells transfected with expression vectors for i) ingetral membrane or ii) secreted WT or variant PAM were labelled for PAM (green), the trans-Golgi network (TGN) (red in i), or insulin (red in ii). DAPI (blue) was used as a nuclear marker. Scale bar, 2μm. Results in C&D are from a single representative experiment that is indicative of at least two independent experiments.

### *PAM* localises to the beta cell secretory pathway

Having established the direction of effect for T2D risk alleles in *PAM*, we next explored a role for PAM in physiologically relevant tissues. Transcript expression profiling detected *PAM* in multiple tissue types, with highest expression in human islets (Figure 1B and Supplementary Figure 2Ai). This is consistent with the published association between *PAM* risk alleles and reduced measures of insulin secretion, and suggests a direct role for PAM in beta cell function. We confirmed broadly similar expression patterns in the publicly available Genotype-Tissue Expression (GTEx) database, though data for pancreatic islets are not available from this project (19). Published RNA sequencing data also showed *PAM* to be equally expressed in enriched fractions of beta cells and non-beta cells (Supplementary Figure 2Aii) (20, 21). We verified this expression pattern at the protein level by immunofluorescent staining of human islets, which revealed co-localisation of endogenous PAM with insulin and glucagon (Figure 1C).

Detailed analysis of the subcellular localisation of WT and variant PAM proteins was then performed for both integral and secreted forms in the human beta cell line EndoC-βH1 (22). Integral membrane WT-PAM immunofluorescence localised to the trans-Golgi network (TGN), where secretory granules originate, whereas secreted WT-PAM displayed a punctate staining pattern and co-localised with endogenous insulin (Figure 1D). Similar results were observed for D563G-PAM and Y651F-PAM (Supplementary Figure 2Bi-ii). As in HEK cells, secreted S539W-PAM expression was undetectable (Figure 1Dii); however, integral membrane S539W-PAM displayed an aberrant staining pattern that localised to the endoplasmic reticulum (ER) as well as the TGN (Figure 1Di and Supplementary Figure 2Biii). These results confirm that PAM localises to components of the beta cell secretory pathway, in agreement with studies in other neuroendocrine cell types (8, 12, 23–25). The S539W substitution may interfere with PAM protein folding such that its translocation from the ER to the TGN is prevented or delayed.

### *PAM* regulates glucose-responsive insulin secretion

Given the negative impact of both T2D risk alleles on overall PAM function, we next determined the effects of reduced PAM levels on beta cell activity. siRNA-mediated knockdown of *PAM* in EndoC-βH1 cells caused a 79% reduction in *PAM* transcript levels (Supplementary Figure 3A), and decreased insulin secretion under both basal (-6.9%, p=0.04) and high glucose conditions (-14.4%, p=8.2x10^-4^) (Figure 2A and Supplementary Figure 4A-B). Furthermore, *PAM* knockdown cells displayed a significantly blunted response to glucose (ratio of high to basal secretion; −7.7 %, p=0.03), indicative of a greater impact on secretion under stimulated conditions. Cellular insulin content was also significantly reduced in *PAM* knockdown cells (-17.6%, p=4.9x10^-5^) despite normal viability and *Insulin (INS)* expression (Figure 2B-D). These results were corroborated by quantitative immunoelectron microscopy, which showed a reduction in the amount of immunogold-labelled insulin per vesicle following *PAM* knockdown (6.26±0.28 vs 8.1±0.41 particles per vesicle (ppv), p=1.77x10^-4^) (Figure 3A&Bi-ii). There were also trends towards decreased cross-sectional vesicle area (0.048±0.003μm^2^ vs 0.056±0.005μm^2^, p=0.18) and increased vesicle density (2.30±0.55 vesicles/μm^2^ vs 1.16±0.09 vesicles/μm^2^, p=0.056) in *PAM* knockdown cells (Figure 3Biii-iv).

**Figure 2.**
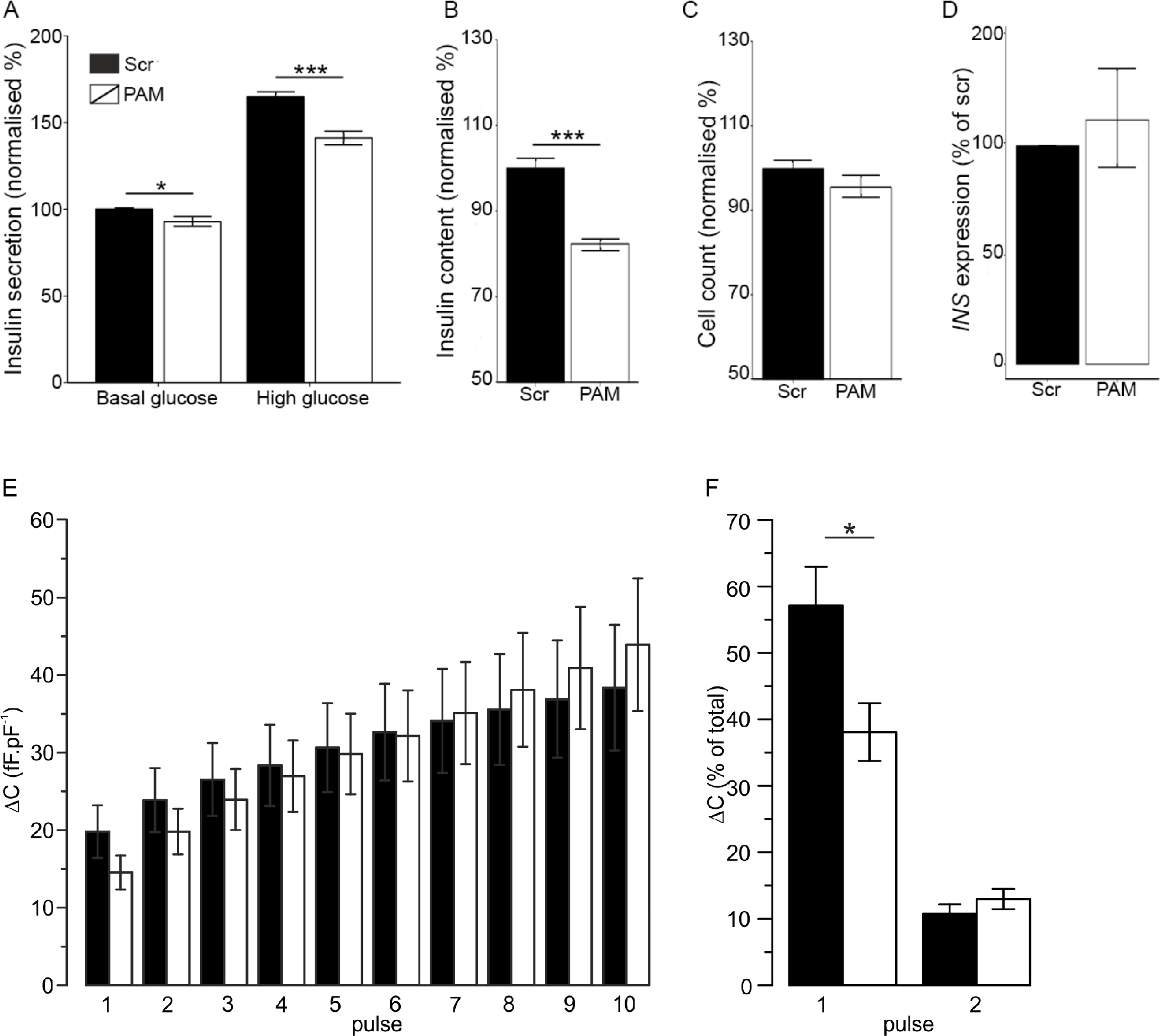
Effects of endogenous PAM on beta cell function. EndoC-βH1 cells were transfected with scrambled (Scr) or *PAM* siRNA then measured for A) insulin secretion (n=8), B) cellular insulin content (n=16), or C) cell viability (n=16), D) *Insulin (INS)* expression (n=3), or E-F) granule exocytosis measured as depolarisation-evoked increased in membrane capacitance (n=15 (Scr) or n=16 (*PAM*)). Error bars are mean ±SEM. P values * <0.05, *** <0.001.

**Figure 3.**
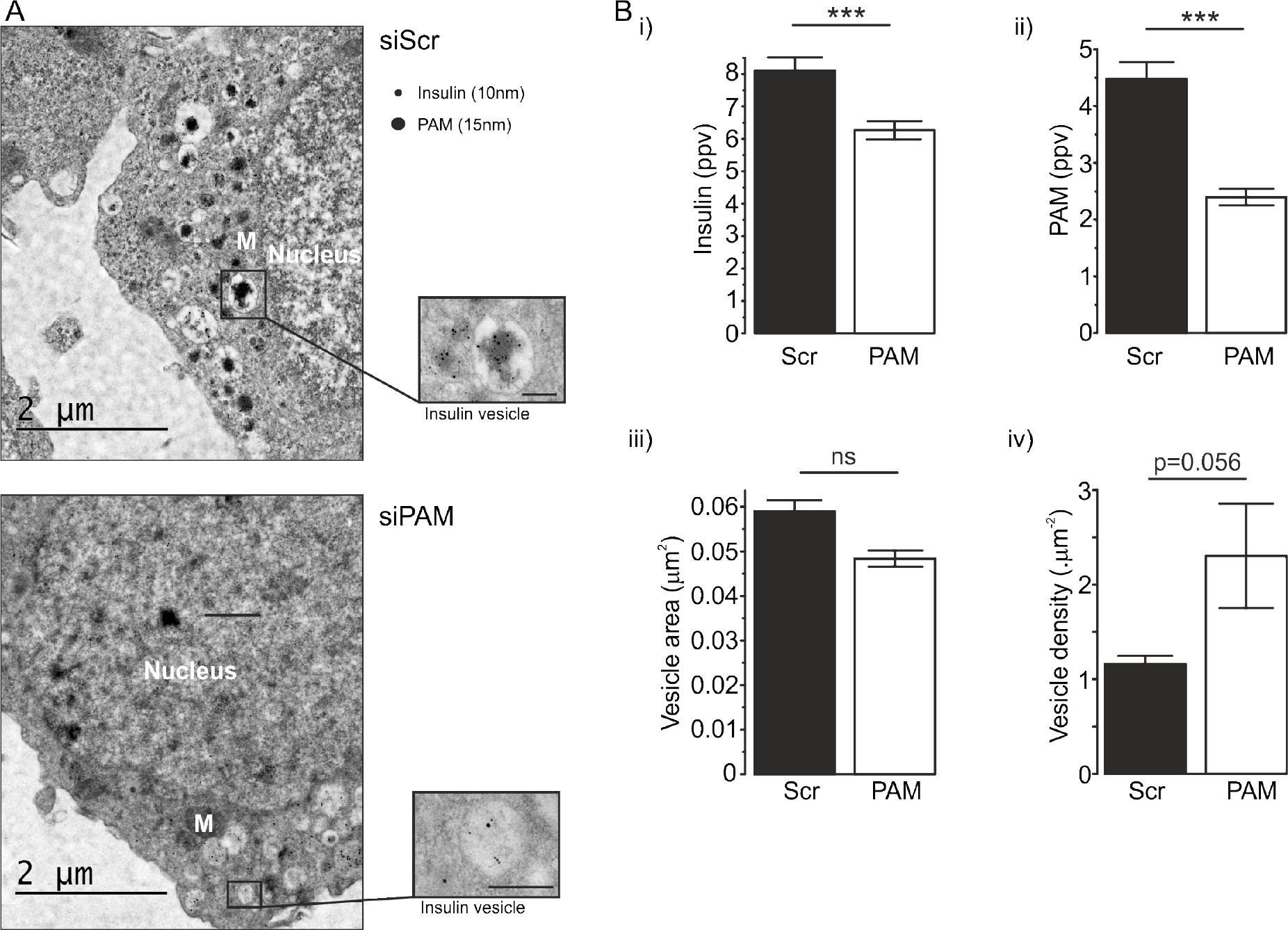
Effects of endogenous PAM on beta cell ultrastructural features. A) Representative electron micrographs from scrambled (Scr) and *PAM* siRNA-treated EndoC-βH1 cells. Scale bar, 2μm. B) i) number of immunogold insulin particles per vesicle (ppv), ii) number of immunogold PAM particles per vesicle, iii) vesicle area, and iv) vesicle density. Results shown are the average of n=235 (Scr) or n=290 (*PAM*) vesicles except for iv) where n=10 cells per treatment. M, mitochondrion. Error bars are mean ±SEM. P value *** <0.001. ns, not significant.

To probe further the effect of *PAM* silencing on insulin secretion, we performed high-resolution single-cell capacitance measurements. Following stimulation of individual cells with a train of 10 depolarisations, an increase in plasma membrane surface area due to vesicle exocytosis could be detected at each step. The cumulative increase in exocytosis was comparable between *PAM* knockdown and control cells (43.9±8.6fF.pF^-1^ vs 38.0±8.1fF.pF^-1^, p=0.64) (figure 2E), and there was no difference in calcium entry (peak current 4.5±0.7pA.pF^-1^ vs 3.6±0.6pA.pF^-1^ for *PAM* knockdown vs control cells, p=0.27) (Supplementary Figure 3B). In contrast, the kinetics of vesicle release were significantly different; in controls cells, nearly 60% of the exocytotic response was elicited by the first pulse, compared with less than 40% in *PAM* knockdown cells (p=0.013) (Figure 2F). The increase in cell capacitance during the first two depolarizations provides an estimate for the size of the readily releasable pool (RRP) of secretory granules, and subsequent pulses represent mobilization of granules from the more intracellular reserve pool (RP) (26). Our results indicate that PAM is an endogenous regulator of insulin availability and the exocytotic response in pancreatic beta cells, with a specific effect on the availability of the RRP.

### Effects of PAM variants on primary beta cell function

Having established a role for PAM in a human beta cell model, we sought to corroborate our results by examining effects for the common *PAM* variant (rs35658696) on primary beta cell function. We analysed exocytosis in single beta cells from dispersed human islets, with capacitance measurements following a train of 10 depolarisations in either 1 or 10 mM glucose. Seven individuals heterozygous for the T2D risk allele were compared to seven age-, gender-, and BMI-matched individuals homozygous for the reference (non-risk) allele (Figure 4A). To specifically test for differences in the RRP, we calculated the total increase in capacitance during the first two depolarisations (Figure 4B). Although no differences were detected at low glucose, a significant decrease in capacitance was seen in risk allele carriers at 10 mM glucose (-26.5%; p = 0.04), indicative of reduced exocytosis from the RRP. Our results are therefore consistent with an effect of rs35658696 status on the kinetics of granule release that manifests predominantly under stimulated conditions.

**Figure 4.**
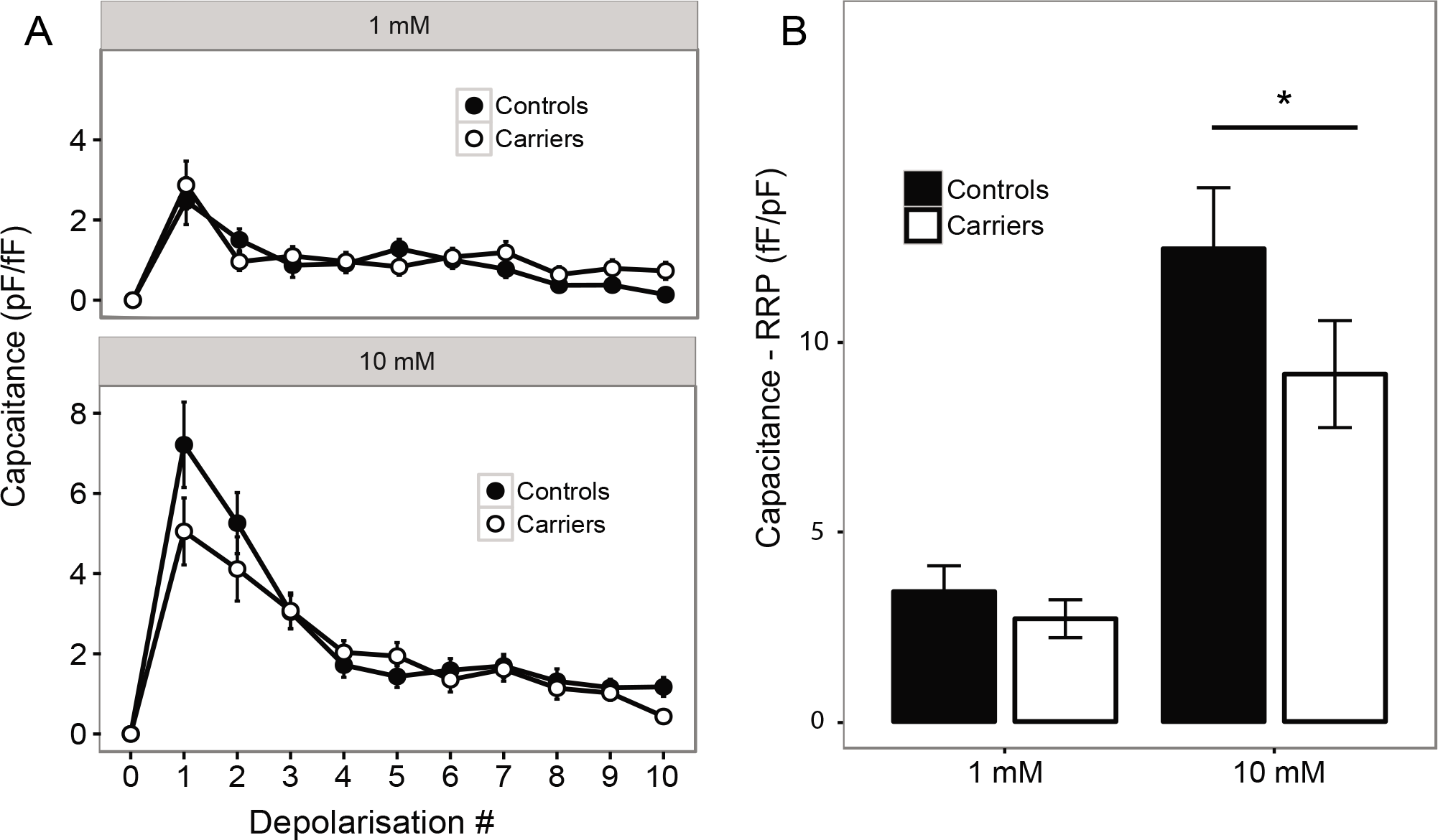
Effects of rs35658696 genotype status on exocytosis measurements in dispersed human islets. Capacitance measurements for primary beta cells from seven individuals heterozygous for the rs35658696 risk allele (“Carriers”) and seven individuals homozygous for the non-risk allele (“Controls”). Dispersed islets were pre-incubated for 1 hour in 1 mM glucose prior to detailed capacitance measurements under the indicated glucose concentrations. (A) Stepwise capacitance measurements after each of ten depolarisations, and (B) the cumulative increase in capacitance for the first two depolarisations, indicative of the size of the RRP. Error bars are mean ±SEM. * P value < 0.05.

### Chromogranin A is a substrate and downstream effector of PAM in beta cells

To elucidate the PAM-regulated pathway in beta cells, we finally sought to identify plausible target proteins that could act as downstream effectors of PAM activity. While any C-terminal glycine-extended peptide could be a substrate for PAM, we reasoned that a key downstream mediator of the observed effects in beta cells would need to be co-expressed with PAM along the secretory pathway. We therefore searched a published catalogue of the islet secretome for potential targets of PAM amidation (27). This highlighted two proteins, Chromogranin A (CgA, encoded by *CHGA*) and Islet Amyloid Polypeptide (IAPP, encoded by *IAPP*), both of which are highly expressed in beta cells and have plausible links with PAM function (27). CgA is a granular packaging protein that self-aggregates to condense large numbers of neuroendocrine peptides into nascent secretory granules (28–30). It can also be cleaved to produce smaller peptides with additional biological roles (31). The role of IAPP in beta cells is less well understood, although deposition of elongated fibrils in islets due to IAPP aggregation is a feature of T2D (32).

*CHGA* silencing (Supplementary Figure 3C) revealed a phenotype similar to that observed for *PAM*, with significant reductions in both insulin content and secretion (Figure 5A-C and Supplementary Figure 4C-D). Using an antibody specific to the glycine-extended (non-amidated) form of CgA (CgA-Gly), we also observed accumulation of CgA-Gly in EndoC-βH1 cells treated with *PAM* siRNA by Western blot analysis (Figure 5D and Supplementary Figure 3D). Treatment of cells with 4-phenyl-3-butenoic acid (4P3BA), a PAM antagonist, produced similar results, demonstrating that amidation of CgA in beta cells is dependent on PAM activity (Figure 5D and Supplementary Figure 3D) (33, 34). An antibody that recognises all forms of CgA (irrespective of amidation status) detected an additional lower molecular weight (∼70kDa) form of CgA, presumably due to cleavage near its C-terminus. Production of this cleaved form was altered in response to decreased PAM expression and activity, suggesting that processing of CgA may be regulated by PAM-dependent amidation (Figure 4D). These results confirm that CgA is a PAM substrate in beta cells, and a strong candidate for mediating the effects of PAM on beta cell function. In contrast, silencing of *IAPP* in EndoC-βH1 cells had no effect on insulin content or secretion (Supplementary Figure 3E-H).

**Figure 5.**
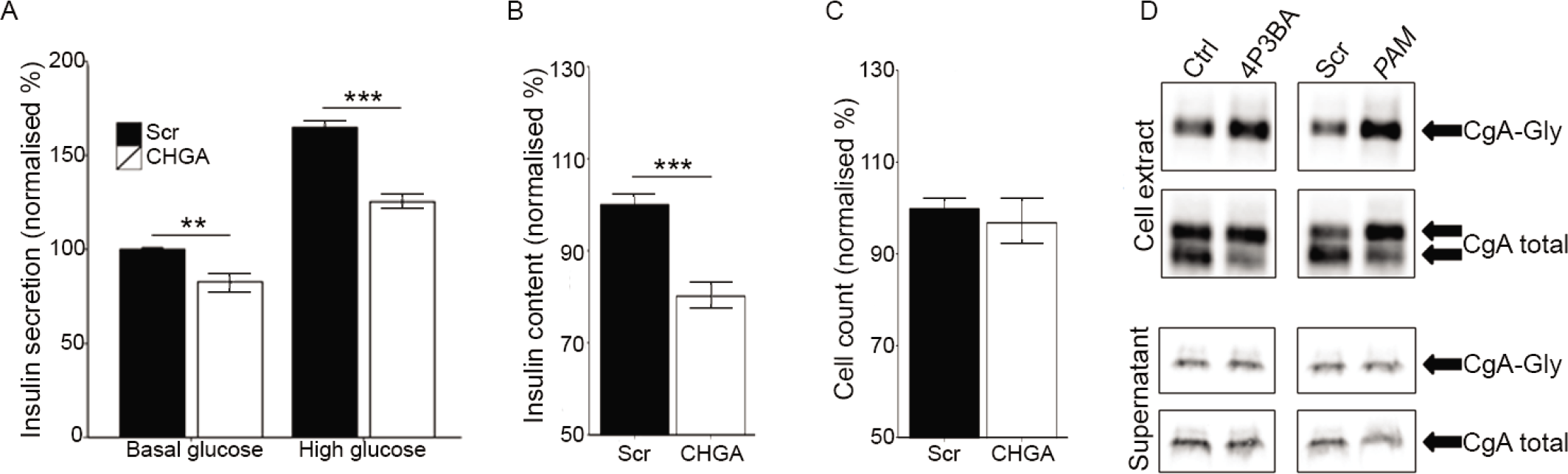
Chromogranin A is a PAM substrate and downstream PAM effector in beta cells. EndoC-βH1 cells were transfected with scrambled (Scr) or *CHGA* siRNA then measured for A) insulin secretion (n=8), B) cellular insulin content (n=16), and C) cell viability (n=16). D) EndoC-βH1 cells were transfected with scrambled or *PAM* siRNA or treated with vehicle (Control) or 4P3BA, then analysed by Western blot using antibodies specific for non-amidated CgA (CgA-Gly) and total CgA. Results in D are from a single representative experiment that is indicative of at least two independent experiments. Error bars are mean ±SEM. P values ** <0.01, *** <0.001

## DISCUSSION

Recent efforts to catalogue coding variants associated with T2D risk have identified independent association signals in the *PAM* gene. The coding risk alleles are also associated with decreased insulinogenic index, a physiological measure of insulin secretion in response to a glucose challenge, indicating a likely endogenous role for PAM in beta cell function. Here, we provide evidence that both T2D-risk alleles reduce overall PAM activity. We also show that reduced PAM expression causes beta cell dysfunction through multiple effects on insulin availability, secretion, and the dynamics of granule exocytosis. Importantly, these defects preferentially affect insulin secretion under stimulated conditions, leading to a blunted glucose response. In line with these results, we observed a consistent effect of the rs35658696 T2D risk allele (p.D563G) on measures of exocytosis in primary human beta cells. Taken together, our findings support a molecular mechanism whereby T2D risk alleles in *PAM* decrease PAM function, directly influencing the ability of pancreatic beta cells to mobilize insulin in response to glucose.

This novel mechanism provides a possible explanation for the relatively larger effect of rs78408340 (p.S539W PAM) that has been reported in two independent GWAS for T2D and insulinogenic index (5, 6). Mislocalisation and/or misexpression of p.S539W PAM would severely affect its function, particularly in cell types where the secreted enzyme predominates. Retention of integral membrane p.S539W PAM within the ER, as indicated by our subcellular localisation studies, could exacerbate this effect through ER stress. The precise cellular dynamics that yield secreted PAM in beta cells have not been elucidated; however, it appears generally to be more predominant in neuroendocrine cell types (15). This raises the intriguing possibility of tissue-specific haploinsufficiency in p.S539W PAM carriers.

To explore the molecular mechanism underlying beta cell dysfunction in *PAM* risk allele carriers, we performed detailed characterisations of the human beta cell line EndoC-βH1 following *PAM* knockdown. The observed defects were broadly similar to those seen in primary human beta cells, with differential effects under stimulated conditions and a significant reduction in the size of the RRP. Interestingly, the effects of reduced PAM function under low glucose conditions in EndoC-βH1 cells were not apparent in primary beta cells. This may reflect differences in the degree of PAM loss-of-function between siRNA-treated EndoC-βH1 cells and rs35658696 risk allele carriers (as suggested by our *in vitro* assay), as well as methodological differences in pre-incubation time prior to capacitance measurements, which could affect the extent of granule repletion.

Our molecular studies in EndoC-βH1 cells confirm that CgA is a substrate for PAM amidation in beta cells (34). CgA is a granular packaging protein known to influence vesicle composition in other neuroendocrine cell types by facilitating the condensation of secreted proteins within nascent secretory vesicles (28–30, 35–37). Previous studies using global *Chga* knockout mouse models have attempted to investigate a role for CgA in pancreatic islets, but have been confounded by compensatory increases in other granin proteins including Chromogranin B (CgB) (38, 39). Our results suggest a mechanism whereby amidation of CgA by PAM enhances its ability to aggregate and package insulin efficiently in granules. In support of this hypothesis, amidation of numerous biological peptides has been shown to promote protein structure and self-aggregation due to improved electrostatic interactions (40–42). C-terminal cleavage of CgA has also been found to increase protein stability in pituitary cells, and the C-terminal domain of PAM interacts with proteins that influence actin cytoskeleton remodelling in yeast-2-hybrid analyses (43–45). These findings provide insights into the mechanism by which PAM and CgA may influence granule biogenesis.

Although our tissue expression analyses showed *PAM* to be most highly expressed in pancreatic islets, they also reveal that *PAM* is widely expressed in human tissues. It is therefore possible that *PAM* could influence beta cell function via amidated signals released from other cell types (21, 46). Glucagon-like peptide 1 (GLP-1) is an amidated peptide released from intestinal L-cells in response to a meal that amplifies insulin secretion (47). Moreover, GLP-1 (7-36-amide) has an increased half-life compared to GLP-1 *in vitro* (7–37), suggesting that amidation improves GLP-1 stability (48). PAM may therefore have paracrine and endocrine effects on beta cell function via decreased GLP-1 amidation and/or secretion, which has implications for genotype-specific responses to incretin-based therapies.

In conclusion, we show that the amidating enzyme PAM is a critical component of the regulated secretory pathway in beta cells, consistent with the observed effects of *PAM* missense variants on T2D risk and insulinogenic index. Our results establish molecular mechanisms for T2D-associated *PAM* risk alleles, and reveal multiple effects of reduced PAM function on insulin granule packaging and release from beta cells, potentially via amidation of CgA. Our study illustrates the importance of coding variants implicated in disease risk to the identification of causal genes for mechanistic studies. Further work is required to explore the possible contribution of other tissues to PAM function, particularly those that reveal implications for incretin-based therapies.

## MATERIALS AND METHODS

### Cloning of PAM expression plasmids

A pCMV6 expression vector encoding integral membrane H. sapiens *PAM* (NM_138821.2) with an in-frame C-terminal Myc-Flag tag (PAM-Myc-Flag) was purchased from Origene. Soluble functional PAM-Myc-Flag encompassing the first 710 amino acids was generated via PCR using primers 5’GTACCCAGAATGTAGCTCC3’ and 5’CGGACGCGTAACTGATCGATGTTCCAA3’. Puromycin-resistant PAM-Myc-Flag expression vectors were generated via *BglII/XmaI* digestion, blunt ending with T4 polymerase, and subcloning into *Hpa*I-digested pIRES puro 2 (Clontech). Mutations were introduced via site-directed mutagenesis using the Stratagene QuikChange II kit (Agilent Biotechnologies) according to manufacturer’s instructions. Primer sequences were 5’CATGTCTGGGATGGAAACTGGTTTGACAGCAAGTTTGTTTAC3’ and 5’GTAAACAAACTTGCTGTCAAACCAGTTTCCATCCCAGACATG3’ for S539W, 5’GACACTATTCTTGTCATAGGTCCAAATAATGCTGCAGTAC3’ and 5’GTACTGCAGCATTATTTGGACCTATGACAAGAATAGTGTC3’ for D563G, and 5’CATTTATGTATCAGATGGTTTCTGCAACAGCAGGATTGTGC3’ and 5’GCACAATCCTGCTGTTGCAGAAACCATCTGATACATAAATG3’ for Y651F. All plasmids were verified by sequencing.

### Cell culture and transfection

Human islets were freshly isolated at the Oxford Centre for Islet Transplantation (OXCIT) in Oxford, UK, as described (49) and cultured in CMRL medium. HEK 293 cells were routinely passaged in DMEM 6429 (Sigma-Aldrich) supplemented with 10% foetal calf serum, 50 units/mL penicillin and 50μg/mL streptomycin. EndoC-βH1 cells were routinely passaged in growth medium supplemented with 5.5mM glucose (22). All cells were maintained at 37°C/5% CO_2_.

For generation of PAM stable cell lines, HEK 293 cells were transfected using Fugene 6 (Promega) according to manufacturer’s instructions. Cells were routinely passaged and maintained in 1μg/mL puromycin. For fluorescence microscopy, EndoC-βH1 cells were transfected in chamber slides (BD Biosciences) using Fugene 6. For siRNA experiments, EndoC-βH1 cells were seeded in Opti-MEM reduced serum medium (Life Technologies) containing 25nM ON-TARGETplus SMARTpool siRNA complexes (Dharmacon) preformed with Lipofectamine RNAiMAX (Life Technologies) according to manufacturer’s instructions. Growth medium was replaced 24hr later and incubated for a further 48hr. For PAM inhibitor experiments, seeded cells were treated with 500μm 4-phenyl-3-butenoic acid (4P3BA) or DMSO vehicle (Sigma-Aldrich) for 72hr.

### Recombinant *PAM* protein production

HEK 293 stable cell lines were seeded into Triple Flasks (Nunc). Medium was replaced with DMEM 1145 supplemented with 1mM sodium pyruvate and incubated for 3-5 days. Supernatant containing secreted recombinant PAM protein was harvested directly from flasks, filtered into a sterile storage bottle (Corning), and pH corrected to 5.5 with HCl. Filtered supernatant was concentrated using a Centricon Plus-70 30kDa MW cut-off Centrifugal Filter Device (Millipore) according to manufacturer’s instructions.

### Quantitation of recombinant *PAM*

For relative quantitation of recombinant PAM proteins, 3x1 μL aliquots of concentrated supernatant were separated on a 4–20% Criterion TGX Stain-Free Gel (Bio-Rad) and transferred to PVDF using a Trans-blot Turbo Transfer Pack (Bio-Rad) according to manufacturer’s instructions. Membranes were incubated overnight in 4A6 anti-Myc tag antibody (Millipore) at 4°C and developed using the Clarity Western ECL Substrate (Bio-Rad) according to manufacturer’s instructions. Proteins were visualised on a ChemiDoc MP (Bio-Rad). Lanes and bands were drawn in Image Lab 5.0 and the average band volume (intensity) quantified for each triplicate of samples. The amount of recombinant variant PAM protein/μL could then be calculated relative to WT-PAM.

### Kinetic analysis of PAM amidating activity

Amidating activity was measured via spectrophotometric detection of chemically converted glyoxylate (18). An assay mix of 100mM MES pH 5.5, 2μM CuSO_4_, 10mM L-ascorbic acid, 0.1mg/mL bovine liver catalase, and 20mM hippuric acid (Sigma-Aldrich) was split into glass vials. Equimolar amounts of recombinant PAM were added to each vial and incubated in a 37°C water bath. Every 30min, samples were removed to a conical 96-well plate (4titude) containing 40mM EDTA pH 8.0. The glyoxylate produced was converted to glyoxylate phenylhydrazone via addition of 0.1% phenylhydrazine HCl (Sigma-Aldrich), incubated at room temperature for 15min, and saturated with concentrated (37%) HCl. The contents of the entire plate were then aspirated and dispensed into a flat-bottomed 96-well plate (Corning) containing 0.2% K_3_[Fe(CN)_6_] using an i-Pipette Automated Pipettor (Apricot Designs). Absorbance of 1,5-diphenylformazan was measured at 515nm using a VersaMax microplate reader (Molecular Devices).

### Kinetic analysis of variant *PAM* substrate affinity

For K_m_ calculations, a 12-point two-fold serial dilution of hippuric acid in which the maximum concentration was 20mM was prepared in assay mix (100mM MES pH 5.5, 2μM CuSO_4_, 10mM L-ascorbic acid, 0.1mg/mL bovine liver catalase). Recombinant PAM or empty vector control supernatant was then added. The amount of D563G-PAM added to the assay was increased such that it displayed equal V_max_ to WT-PAM. The volume of empty vector supernatant matched the volume of WT-PAM used in that experiment. Vials were incubated in a 37°C water bath for 1hr. Samples were then removed into a conical 96-well plate containing 40mM EDTA pH 8.0.

### Gene expression analyses

Gene expression was measured via Taqman assay. RNA was purified from TRIzol (Ambion) homogenates or sourced from commercially available RNA tissue panels (Human Total RNA Master Panel II (Clontech), FirstChoice Human Total RNA Survey Panel (Ambion), and mouse Total RNA Master Panel (Clontech)). Pooled human adult pancreas RNA from n=3 donors and pooled human adult islet RNA from n=3 donors was sourced from the Oxford Islet Biobank (50). Pooled mouse islet RNA was sourced from four 12–28 week-old NMRI mice. Hypothalamus and pituitary RNA were commercially sourced from AMSBio. cDNA was generated using the GoScript Reverse Transcription System (Promega) according to manufacturer’s instructions and analysed using an Applied Biosystems 7900HT. C_T_ values were normalised to the combined average of the housekeeping genes *PPIA* and *TBP. PAM* expression in human islets and sorted beta cells and non-beta cells was analysed based on previously published RNA sequencing data (20).

### Immunofluorescent labelling and microscopy

For light microscopy, analyses were performed on 4μm sections. Human pancreatic specimens fixed in 10% formalin and embedded in wax were used to localise PAM in islets. Sections were de-waxed, rehydrated, and heated to 95°C in 10mmol/L Tris + 1mmol/L EDTA pH 6.0 for 10min for antigen retrieval. Tissue sections were incubated overnight in anti-PAM (Abcam ab109175), guinea-pig anti-insulin (in-house antibody), and mouse anti-glucagon (Sigma-Aldrich G2654) antibodies. PAM immunoreactivity was enhanced using a tyramide amplification step (Invitrogen) and visualised with Alexa Fluor 488 (Life Technologies). Insulin and glucagon were visualised using Alexa Fluor 633 (Life Technologies) and anti-mouse TRITC (Sigma-Aldrich) respectively.

EndoC-βH1 cells transfected with *PAM* expression vectors were fixed in 4% paraformaldehyde, permeabilised in 0.1% (v/v) Triton X-100, and blocked in 10% (w/v) BSA. Primary antibodies (4A6 anti-Myc, Belfast anti-insulin, TGN46 anti-trans Golgi [Sigma-Aldrich], and H-70 anti-calnexin [Santa Cruz Biotechnology sc-11397]) were incubated overnight at 4°C. Slides were then incubated in Alexa Fluor-conjugated secondary antibodies (Life Technologies) and mounted in Vectashield Mounting Medium with DAPI (Vector Biolaboratories). Cells were visualised using an LSM 10 META confocal laser scanning module arranged on an Axiovert 200 microscope and a Plan-Apochromat 63x/1.4 oil immersion objective (Carl Zeiss). An argon laser was used to excite Alexa Fluor 488 at λ=488nm and a HeNe laser was used to excite Alexa Fluor 543 at λ=543nm. DAPI was excited in two-photon mode using the 740nm line of an infrared light Chameleon laser.

### Insulin secretion and membrane capacitance assays in EndoC-βH1

Insulin secretion assays were performed as previously described (51). The CyQUANT Direct Cell Proliferation Assay (Thermo Fisher) was used to count cells according to manufacturer’s instructions. Insulin secretion and content measurements were normalised to cell count on a per-well basis to account for differences in the number of plated cells. Corrected values were then normalised to basal insulin secretion rates for scrambled on a per-experiment basis to eliminate variability in baseline secretion rates across biological replicates. Electrophysiological measurements were performed using an EPC10 Patch Clamp Amplifier (HEKA) at 32°C using the standard whole cell configuration as previously described (52). The pipette solution (intracellular medium) contained 125mM Cs-glutamate, 20mM CsCl, 15mM NaCl, 1mM MgCl_2_, 5mM HEPES, 0.05mM EGTA, 0.1mM cAMP, and 3mM Mg-ATP (pH 7.15 with CsOH). The extracellular solution contained 1mM glucose, 118mM NaCl, 5.6mM KCl, 1.2mM MgCl_2_, 5mM HEPES, and 2.6mM CaCl_2_ (pH 7.4 with NaOH). Tetraethylammonium chloride (20mM) was added to block residual K^+^ currents.

### Electron microscopy

Transfected EndoC-βH1 cells were fixed in 2.5% glutaraldehyde, post-fixed in 2% uranyl acetate, dehydrated in graded methanol, and embedded in London Resin Gold (Agar Scientific). Ultrathin sections cut onto nickel grids were immunolabelled for PAM (Abcam ab109175) followed by anti-rabbit biotin (Vector Laboratories) and streptavidin gold 15nm (Biocell). Insulin was immunolabelled using an in-house antibody followed by anti-guinea pig gold 10nm (Biocell). Sections were viewed on a Joel 1010 microscope (accelerating voltage 80kV) with a digital camera (Gatan). To estimate PAM concentration in secretory vesicles, semi-automatic quantification of immunogold labelling was performed by a blinded observer using Image J software. The following parameters were determined: cross-sectional vesicle area, vesicle density (number of vesicles/cytoplasmic area), and insulin or PAM labelling/vesicle (number of immunogold particles/vesicle).

### Western blot analysis of *CgA* amidation

Whole-cell extracts (WCE) from siRNA or inhibitor-treated EndoC-βH1 cells were prepared as previously described (53). Supernatant was harvested directly from cells and filtered with a 0.22μm filter. Protein yields were determined by the Bio-Rad Protein Assay according to manufacturer’s instructions. We routinely analysed 2-10μg WCE and 0.5% total supernatant volume via denaturing SDS-PAGE. Antibodies specific for CgA-Gly (Abcam ab52983) (34) and total CgA (Abcam ab15160) were used to calculate the ratio of amidated to non-amidated CgA using Image Lab 5.0.

### Analysis of exocytosis in primary human beta cells

Human pancreatic islets were isolated and prepared at the Alberta Diabetes Institute IsletCore as previously described (54). Single-cell capacitance responses were monitored as described above, with an additional pre-incubation period of 1 hour at 1 mM glucose. Following recordings, cells were positively identified by insulin immunostaining and islet samples were genotyped as previously described (50). We examined the kinetics of exocytosis at 1 and 10 mM glucose for seven risk allele carriers and seven controls, matched as groups for age (54.9 years [carriers] vs. 58.4 years [controls]; p = 0.52), gender (2/7 females [carriers] vs. 4/7 females [controls]) and BMI (27.7 [carriers] vs. 26.7 [controls]; p = 0.68). Mean capacitance was calculated per condition for each depolarisation step, and ANOVA with Tukey’s post-hoc test was applied to detect statistically significant differences.

### Statistics

All statistical analyses were performed in R v3.0.2, except electron microscopy data and electrophysiological measurements in EndoC-βH1 cells, which were analysed using OriginPro v8.5.1. Graphs show means of the indicated number of replicates, and error bars are SEM. For kinetic data, rates were found to be dependent on substrate concentration, and exponential models were fitted. Kinetic and cellular assay data were analysed using two-sided Welch’s t-test; electrophysiological, electron microscopy, and Western data using paired student’s t-test.

### Study Approval

All studies were approved by the University of Oxford’s Tropical Research Ethics Committee (OxTREC reference 2–15), or the Oxfordshire Regional Ethics Committee B (REC reference 09/H0605/2). Human pancreatic tissue was obtained from subjects at post-mortem examination according to local and national ethics permissions.

## ACKNOWLEDGEMENTS

The authors acknowledge sharing of data from the GoT2D and T2D-GENES consortia prior to publication. We thank Dr Juris Galvanovskis, University of Oxford, for microscopy assistance, and James Lyon at the Alberta Diabetes Institute IsletCore for his work on human islet isolations. We also thank the Human Organ Procurement and Exchange Program (Edmonton) and the Trillium Gift of Life Network (Toronto) and other organ procurement agencies for their efforts in obtaining human pancreases for research.

## Author contributions

A.R. S.K.T. B.H. M.M.U. N.L.B. C.J.G. A.C. P.E.M. and A.L.G. designed experiments, A.R. S.K.T. B.H. M.M.U. A.B. and A.C. performed experiments, A.R. S.K.T. B.H. M.M.U. N.L.B. C.J.G. A.C. P.E.M. P.R. and A.L.G. interpreted data, A.R. S.K.T. and A.L.G. wrote the manuscript. All authors approved the final draft of the manuscript.

